# Mediation Analysis with Compositional Exposures

**DOI:** 10.64898/2026.09.24.754147

**Authors:** Jing Ma, Poorbita Kundu, Timothy Randolph, Sandi Navarro, Meredith Hullar

## Abstract

Understanding the causal pathways linking the gut microbiome to downstream biomarkers and clinical outcomes is central to microbiome research. When the microbiome acts as the exposure, the high dimensionality and compositional structure of the data introduce unique methodological challenges for mediation analysis. To address these challenges, we propose a latent variable mediation framework that captures variation in microbial composition through a microbial balance, defined as the log-ratio between two unknown subsets of taxa. This balance serves as a latent scalar exposure and simplifies the estimation of the overall indirect effect at the community level, while simultaneously identifying specific taxa that contribute to the overall direct and indirect effects. We describe the model’s estimation and inference, and illustrate the method using data from the Multiethnic Cohort–Adiposity Phenotype Study, where we examine the mediation pathway from the gut microbiome through lipopolysaccharide-binding protein, a marker of metabolic endotoxemia, to percent liver fat.

## 1 Introduction

Microbial-derived products are key mediators of host-microbiome interactions (Roager and Licht, 2018; Glick et al., 2024). These include microbial *metabolites*, such as short-chain fatty acids and indole derivatives (Mendis et al., 2025; Zhang et al., 2021; Su et al., 2022), and structural *cell-envelope components* such as lipopolysaccharide (endotoxin), which upon translocation across the gut barrier triggers low-grade inflammation (Cani et al., 2007). In some settings the mediator of interest is not the microbial product itself but a circulating host marker of microbial exposure that it induces (Moreno-Navarrete et al., 2012). Delineating how these products transmit the effect of the gut microbiome on host outcomes would help identify intervention targets and clarify the mechanisms of microbiome-associated disease. Yet this pathway—with the microbiome as a compositional *exposure*, a microbial product or its host marker as the mediator, and a clinical endpoint as the outcome—remains understudied.

Modeling the microbiome as an exposure raises distinct challenges. First, the data are compositional: abundances carry only relative information (Gloor et al., 2017), and are further complicated by overdispersion and excess zeros (Martín-Fernández et al., 2011). Modeling each taxon independently from other taxa invites spurious associations (Li et al., 2024, 2025). Second, a microbial composition is high-dimensional, and no log-ratio representation is simultaneously full-rank, interpretable, and free of arbitrary choices: the centered log-ratio (CLR) is rank-deficient, so its coordinates are collinear and their coefficients are not separately identified, the isometric log-ratio (ILR) transform (Egozcue et al., 2003) requires a basis that depends on the ordering of taxa and yields coordinates without direct biological meaning, whereas the additive log-ratio (ALR) transform (Aitchison, 1986) requires a reference taxon that often lacks clear justification. Third, and most crucially, the effect of a high-dimensional exposure admits no single interventional contrast (Kramer et al., 2026): direct and indirect effects become well-defined only after the exposures are reduced to low-dimensional representations. Such a reduction must be *supervised* rather than performed without reference to the mediator and outcome (Nabi et al., 2017), as in unsupervised principal component analysis (PCA), whose leading components need not capture the mediating signal (Zhao et al., 2022). The reduction must also remain interpretable at the level of individual taxa (Pawlowsky-Glahn et al., 2011), since these taxa specify the intervention whose effect is estimated. Existing summaries of high-dimensional exposures, such as principal-component projections (Zhao, 2024; Nabi et al., 2017) or weighted indices (Carrico et al., 2015), are not built for compositional data and respect neither the simplex constraint nor taxon-level interpretability.

To date, compositional mediation analysis has been developed almost entirely for the complementary setting in which the microbiome is the *mediator*. For example, Sohn and Li (2019) and Sohn et al. (2022) used simplicial algebra (Aitchison et al., 2002) to model the compositional mediator; Wang et al. (2020) modeled taxa with a Dirichlet distribution, though it can be too restrictive for overdispersed microbiome counts; Fu et al. (2023) extended Wang et al. (2020) with a Dirichlet-multinomial model, creating uncorrelated mediators through the ILR transformation at the cost of an ordering-dependent, hard-to-interpret representation; and Hong et al. (2023) studied mediators in the form of summated log-ratio (Greenacre, 2020) defined at internal nodes of a phylogenetic tree. Parallel to these works that study the relative mediation effects of individual taxa, Huang (2020) introduced a latent-variable model based on balances, defined as log ratios between *two groups of taxa* (Egozcue and Pawlowsky-Glahn, 2005), for simultaneous inference on the presence of any relative mediation effects. Balances give an interpretable, scale-invariant representation whose numerator and denominator can be read as pathogenic versus commensal taxa (Rivera-Pinto et al., 2018) and linked to metabolic functions or ecological guilds (Wu et al., 2021). Despite these advancements, *no rigorous framework models microbiome compositions as exposures in mediation*.

A microbial balance is precisely the summary needed for a compositional exposure: a single log-ratio between two groups of taxa, reducing a high-dimensional composition to one scalar while naming the taxa that define it. In this paper we develop a mediation framework built on such balances, used as latent exposures that jointly drive the mediator and the outcome, so that both the direct and indirect effects are simple functions of a single log-ratio. Unlike PCA, which is estimated separately from the outcome, the balance is learned *jointly* with the other model parameters, aligning the dimensionality reduction with the causal inference task while selecting the contributing taxa. We develop a Bayesian inference procedure that estimates the balance configuration and the mediation effects together using Gibbs and Metropolis–Hastings sampling (Tierney, 1994; Hastings, 1970). Through simulation studies, we evaluate the method’s accuracy in estimating the indirect effect and in recovering the taxa that define the balance, including its robustness to model misspecification and to unmeasured mediator–outcome confounding. We then illustrate the method using data from the Multiethnic Cohort–Adiposity Phenotype Study (Hullar et al., 2021, MEC-APS), examining whether lipopolysaccharide-binding protein, a circulating marker of microbial endotoxin exposure, mediates the effect of the gut microbiome on percent liver fat. The rest of the paper is organized as follows. Section 2 introduces the balance-based structural equation model and defines the direct and indirect effects. Section 3 develops the Bayesian inference procedure. Section 4 presents simulation studies, and Section 5 applies the method to the MEC-APS study. We conclude with a discussion in Section 6.

## 2 Models and Estimands

Consider *n* subjects, each with a *d*-dimensional compositional exposure vector ***x***_*i*_ = (*x*_*i*,1_, …, *x*_*i,d*_)^T^ where 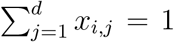. Let *m*_*i*_ *∈ ℝ* denote a scalar mediator, ***c***_*i*_ = (*c*_*i*,1_, …, *c*_*i,k*_)^T^ *∈ℝ*^*k*^ the *k*-dimensional vector of covariates, and *y*_*i*_ *∈ ℝ* a scalar continuous outcome. Our scientific goal is to investigate how the compositional exposure ***x***_*i*_ influences the outcome *y*_*i*_ both directly and indirectly through the mediator *m*_*i*_, for *i* = 1, …, *n*. For clarity of exposition, we develop the framework for a single mediator; extensions to multiple, possibly correlated, mediators are discussed in Section 6.

### 2.1 Structural Equation Model

We assume there exists a scalar latent variable *B* (***z***, *X*), defined as a weighted log-ratio between two subsets of variables in a composition *X* (Egozcue and Pawlowsky-Glahn, 2005), that has direct effects on both the mediator and the outcome (Figure 1). Here ***z*** = (*z*_1_, …, *z*_*d*_) is a *d*-dimensional indicator vector assuming values 1, −1, and 0, where *z*_*j*_ = 1 indicates that part *j* of the composition is in the numerator of the balance, *z*_*j*_ = *−*1 indicates the denominator, and *z*_*j*_ = 0 indicates that the part is not a member of the balance. Let *Ƶ*_+_, *Z*_*−*_ and *Ƶ*_0_ be non-overlapping sets of indices of ***Ƶ***, and *Ƶ*_+_ *∪ Ƶ*_*−*_ *∪ Ƶ*_0_ = {1, …, *d*}. Given ***z***, the balance is defined as

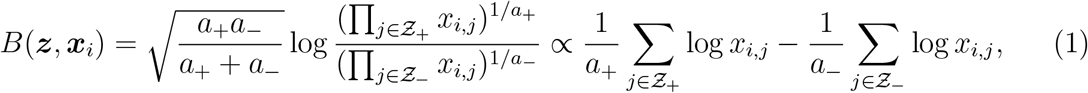

**Figure 1.**
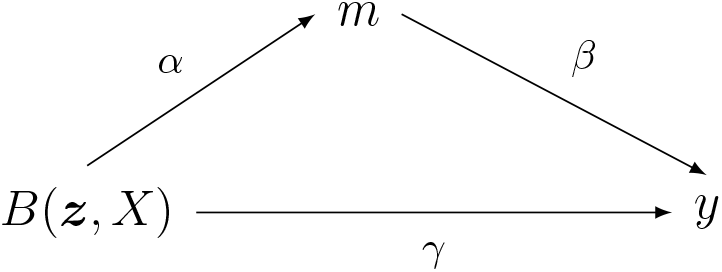
Mediation paths from a latent balance *B*(***z***, *X*) to a scalar mediator *m* and a scalar outcome *y*.

where *a*_+_ and *a*_*−*_ denote the number of indices in *Ƶ*_+_ and *Ƶ*_*−*_, respectively, and the sign *∝* means proportional to.

The balance in (1) is defined for strictly positive relative abundances. Microbiome data often contain many zeros, which should be imputed using pseudocounts, as done in the MEC application, or model-based approaches (Martín-Fernández et al., 2015; Zhou et al., 2025; Luo et al., 2025) before being passed to the proposed method.

The key of our proposal is to associate the observed compositions ***x***_*i*_, the mediator *m*_*i*_, and the outcome *y*_*i*_ via the unobserved balance configuration vector ***z***. Specifically, we study the path relations via the following linear structural equation model (SEM)

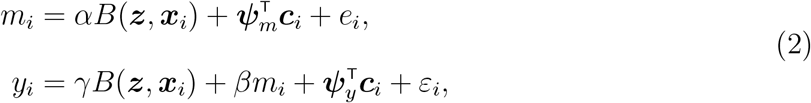

where the error terms 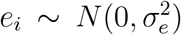 and 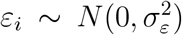. Here ***ψ***_*m*_ *∈ ℝ* ^*k*^ and ***ψ***_*y*_ *∈ℝ*^*k*^ denote the covariate coefficient vectors in the mediator and outcome regression models, respectively. Our primary goal is to estimate the model parameters *α, β, γ*, ***ψ***_*m*_, ***ψ***_*y*_ and ***z*** given observations 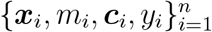.

### 2.2 Identifiability and sign convention

The balance in (1) is invariant to permutations of the taxa within *Ƶ*_+_ and within *Ƶ*_*−*_, but exchanging the numerator and denominator of the balance changes its sign, *B*(*−****z, x***_*i*_) = *−B*(***z, x***_*i*_). Accordingly, replacing (***z***, *α, γ*) with (*−****z***, *−α, −γ*) leaves the model (2) unchanged. To resolve this sign ambiguity, we impose the constraint *γ >* 0, so *Ƶ*_+_ denotes the group of taxa whose higher abundance, relative to *Ƶ*_*−*_, is associated with a higher outcome through the direct path. The normalizing constant in (1) fixes the scale of the balance. Conditional on ***z***, the remaining parameters in (2) are identified under the usual rank conditions for linear regression, provided that both *Ƶ*_+_ and *Ƶ*_*−*_ are non-empty and *B*(***z, x***_*i*_) is not collinear with the covariates ***c***_*i*_.

### 2.3 Identification of direct and indirect effects

For a contrast in the latent balance from *b*^*∗*^ to *b*, the linear SEM in (2) yields the natural direct effect

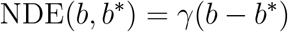

and the natural indirect effect

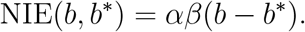

Here, the contrast *b*^*∗*^ *→ b* represents a change in the underlying composition that changes the relative abundance of taxa in *Ƶ*_+_ versus *Ƶ*_*−*_ from balance value *b*^*∗*^ to *b*; compositions yielding the same balance value define the same exposure level under the model. Thus, for a one-unit increase in the balance, the direct, indirect, and total effects are *γ, αβ*, and *γ* + *αβ*, respectively.

These effects are identified under the standard assumptions for causal mediation analysis: the stable unit treatment value assumption, positivity, and no unmeasured exposure– outcome, exposure–mediator, or mediator–outcome confounding conditional on ***c***_*i*_ (Rubin, 1980; Rosenbaum and Rubin, 1983; VanderWeele, 2016). Identification further requires that no mediator–outcome confounder are affected by the exposure (VanderWeele et al., 2014). Under these assumptions, together with the linear models in (2) and the absence of an exposure–mediator interaction in the outcome model, the above expressions follow from the mediation formula (Imai et al., 2010; VanderWeele and Vansteelandt, 2014).

## 3 Bayesian Inference

If the balance configuration ***z*** is known, the model in (2) can be estimated by ordinary least squares applied to the mediator and outcome regressions (VanderWeele, 2016). When ***z*** is latent, however, estimating it separately from the two regressions can yield conflicting balance configurations, since the mediator and outcome models need not favor the same ***z***. Instead, we provide a Bayesian solution that jointly estimates ***z*** and the effects.

### 3.1 Priors

We impose the following priors on the coefficients

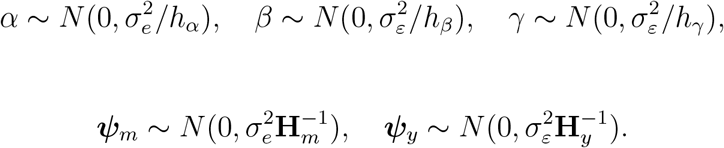

For the variances, we use conjugate inverse-gamma priors 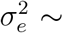 Inverse-Gamma(*ν/*2, *λ*_*e*_/2) and 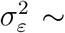 Inverse-Gamma(*ν/*2, *λ*_*ε*_/2). The advantage of the normal-inverse-gamma priors is that the coefficients and variances can be integrated out in closed form. Here we treat *h*_*α*_, *h*_*β*_, *h*_*γ*_, **H**_*m*_, **H**_*y*_, *ν, λ*_*e*_ and *λ*_*ε*_ as hyperparameters. We place uninformative priors for the coefficients such that *h*_*α*_ = *h*_*β*_ = *h*_*γ*_ = 10^*−*6^ and **H**_*m*_ = **H**_*y*_ = 10^*−*6^**I**_*k*_ with **I**_*k*_ being the *k × k* identity matrix. For the variances we set *ν* = *n, λ*_*e*_ to the sample variance of the mediator, and *λ*_*ε*_ to the sample variance of the outcome. For the balance configuration ***z***, we impose an independent categorical prior on each component,

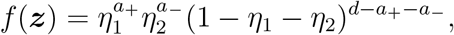

where *η*_1_ and *η*_2_ are the prior probabilities that a taxon belongs to *Ƶ*_+_ and *Ƶ*_*−*_, respectively, so that the expected numbers of parts are *dη*_1_ and *dη*_2_. A non-informative choice is *η*_1_ = *η*_2_ = 1/3, while smaller values lead to more parsimonious selection of taxa.

The model in (2) conditions on the observed relative abundances ***x***_*i*_. To account for sampling variability in compositional counts, the model can be extended by introducing latent proportions and specifying a sampling model for the counts, such as the Dirichlet-multinomial (Chen and Li, 2013) or the logistic-normal multinomial (Silverman et al., 2019). Either extension can be readily incorporated into the Bayesian inference procedure developed here. In this work, however, we treat ***x***_*i*_ as fixed to avoid introducing additional computational complexity beyond that associated with selecting ***z***, as described below. A graphical representation of the proposed mediation framework is provided in **Fig**. 2.

### 3.2 Posterior inference with Gibbs sampler

We begin by introducing several matrix notations to simplify the posterior derivation. Let

***y*** = (*y*_1_, …, *y*_*n*_)^T^, ***m*** = (*m*_1_, …, *m*_*n*_)^T^, **X** = (***x***_1_, …, ***x***_*n*_)^T^ and **C** = (***c***_1_, …, ***c***_*n*_)^T^. Let **B**_*z*_ = (*B*(***z, x***_1_), …, *B*(***z, x***_*n*_))^T^. Let **D**_*m*_ = [**B**_*z*_, **C**] and **D**_*y*_ = [***m*, B**_*z*_, **C**] denote the design matrix in the mediator and outcome regression model, respectively. Conditional on the balance configuration ***z***, the joint likelihood is the product of the likelihood in the outcome regression model 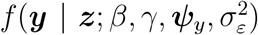 and the likelihood in the mediator regression model 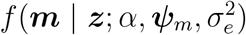. Thanks to the normal-inverse-gamma priors, we can integrate out the coefficients (*α, β, γ*, ***ψ***_*m*_, ***ψ***_*y*_) and the variance parameters 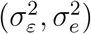 to obtain the conditional density of observed data *f* (***y, m*** | ***z***). Consequently, the conditional distribution of the balance configuration ***z*** given observed data is

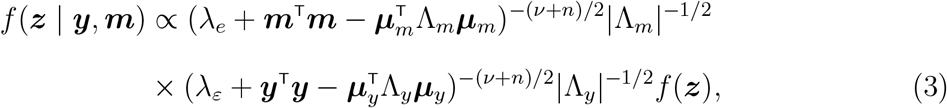

Where 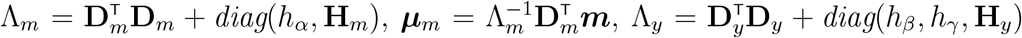 and 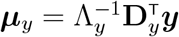.

Because the coefficients and variances have been integrated out in (3), the balance configuration ***z*** can be updated directly from its collapsed conditional, which improves mixing. We update ***z*** one coordinate at a time using a Gibbs sweep. At each iteration we visit the *d* coordinates in a random order; for coordinate *j*, the full conditional of *z*_*j*_ given the remaining entries ***z***_*−j*_ and the data is the three-category distribution

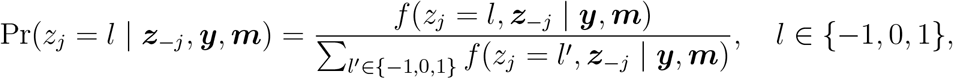

which is available in closed form from (3). We draw *z*_*j*_ exactly from this distribution, restricting its support to values that keep both *Z*_+_ and *Z*_*−*_ nonempty so that the balance remains well-defined.

Conditional on ***z*** and observed data, the posterior of (*α*, ***ψ***_*m*_) after integrating out the variance is a multivariate *t*-distribution with degrees of freedom *ν* + *n*, location parameter ***µ***_*m*_, and shape parameter

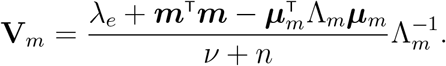

Similarly, the posterior of (*β, γ*, ***ψ***_*y*_) given observed data and ***z*** is a multivariate *t*-distribution with degrees of freedom *ν* + *n*, location parameter ***µ***_*y*_, and shape parameter

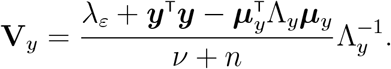

Within each iteration of the MCMC, we recompute the balance **B**_***z***_ after updating ***z*** and sample the coefficients from their conditional posteriors. To enforce the sign convention *γ >* 0 of Section 2.2, we then relabel the draw: if the sampled *γ* is negative, we replace (***z*, B**_***z***_, *α, γ*) by (*−****z***, *−***B**_***z***_, *−α, −γ*), equivalently swapping *Z*_+_ and *Z*_*−*_. Algorithm 1 summarizes the details of one MCMC iteration within the Gibbs sampler; full derivation details are provided in Appendix A. The convergence of the MCMC algorithm can be evaluated using chains with different starting points and by examining the cumulative moving average for each component of ***z***. After convergence, we estimate the component-wise posterior probabilities of the three categories (+1, *−*1, 0) by their marginal proportions across the retained draws; the posterior inclusion probability (PIP) of a taxon is its probability of being nonzero, Pr(*z*_*j*_ *≠*0). Taxon selection is done by thresholding the PIPs at a desired level. Finally, posterior inference for the direct and indirect effects can also be obtained using these MCMC samples.

#### Algorithm 1

One MCMC iteration with a Gibbs update of *z* for compositional mediation analysis

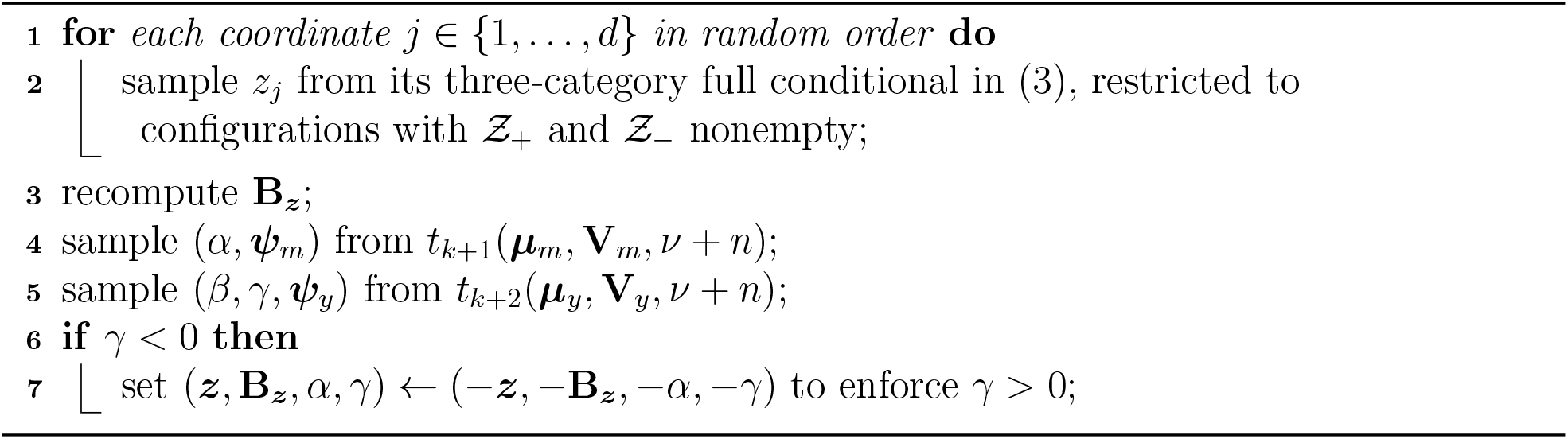

### 3.3 Scaling to high dimensions with Metropolis–Hastings

The Gibbs sweep updates every coordinate of ***z*** at each iteration, which requires evaluating the collapsed density (3) at the three candidate values for each of the *d* coordinates. When *d* is very large, a faster alternative is the Metropolis–Hastings (MH) sampler of Huang (2020), which changes a single coordinate per iteration. Given the current ***z***, it selects a pair of values (*l, l*′) with probability 1/3 and proposes ***z***′ by modifying one eligible coordinate according to a local proposal 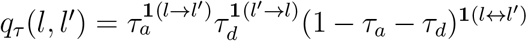, where τ = (*τ*_*a*_, *τ*_*d*_, 1− *τ*_*a*_ − *τ*_*d*_) are the probabilities of value addition, deletion, and switching. The move is accepted with probability

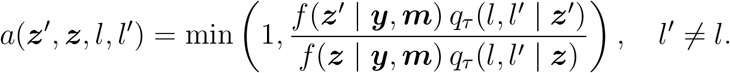

The Gibbs and MH samplers target the same stationary distribution (3), but differ in cost and mixing. The MH sampler evaluates (3) only once per iteration, whereas the Gibbs sweep evaluates it of order *d* times. However, the MH proposal modifies at most one coordinate and can mix slowly when the posterior concentrates on a sparse set of configurations: most proposals are then rejected, and we have observed acceptance rates below 5% in our empirical studies. Mixing can be improved by an informed proposal that preferentially selects coordinates whose marginal inclusion probabilities suggest a likely change (Peixoto, 2025), at the cost of a more elaborate proposal. An empirical comparison of the two algorithms is presented in Section 4.

## 4 Numerical Experiments

### 4.1 Data generation

To generate realistic microbiome data, we sampled zero-inflated count data by rounding continuous data generated from a latent log-normal distribution to the nearest integer. Specifically, we first sampled continuous data from a multivariate normal distribution with mean zero and covariance Σ = (*σ*_*j,j*_*′*) *∈* ℝ^50*×*50^ where *σ*_*j,j*_*′* = 9·0.2^|*j−j*^*′*^|^. Count data were generated by exponentiating the above matrix and rounding. At sample size *n* = 100, about 40% of the resulting count data matrix are zero, mimicking the common sparsity pattern in real data. Finally, zeros in the count data were replaced with 0.5 and a total sum normalization was used to obtain the compositional data 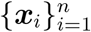. A one-dimensional covariate *c*_*i*_ and the error terms *e*_*i*_ and *ε*_*i*_ were independently sampled from a standard normal distribution, and the covariate coefficients were set to ***ψ***_*m*_ = ***ψ***_*y*_ = 1. The path coefficients are *α* = *γ* = 1/3 and *β ∈ {*0, 2*}*, so that the indirect effect *αβ* equals 0 and 2/3, respectively, and the direct effect *γ* = 1/3.

To generate the mediator and the outcome, we consider scenario I, a model correctly specified according to (2) where ***z*** = (1, 1, 1, *−*1, *−*1, *−*1, 0, …, 0) with equal weights, and scenario II, a model misspecified such that the balance *B*(***z, x***_*i*_) in (2) is replaced by the weighted log contrast log *x*_*i*,1_+0.4 log *x*_*i*,2_+1.2 log *x*_*i*,3_*−*1.5 log *x*_*i*,4_*−*0.8 log *x*_*i*,5_*−*0.3 log *x*_*i*,6_. To ensure that any difference in performance reflects misspecification of the balance *shape* rather than a change in signal strength, we rescale the log contrast in scenario II so that its variance across subjects matches that of the equal-weight balance in scenario I; the path coefficients, and hence the indirect and direct effects, are then identical across the two scenarios. We also consider scenario III, in which a hidden covariate *u*_*i*_ *~ N* (0, 1) affects both the mediator and outcome, with additive contributions *δ*_*m*_*u*_*i*_ and *δ*_*y*_*u*_*i*_, respectively, but is omitted when fitting the model. Scenario III evaluates the impact of unmeasured mediator–outcome confounding, whose induced bias in the estimated indirect effect grows with the product *δ*_*m*_*δ*_*y*_ and vanishes when either loading is zero.

We consider two sample sizes *n ∈ {*100, 200*}* for scenarios I and II, and only *n* = 100 for scenario III and all sensitivity settings. For each fit we ran the Gibbs sampler for 2 *×* 10^4^ iterations with a burn-in of 10^4^. Under scenario I, *β* = 2 and *n* = 100, we additionally ran the Metropolis–Hastings algorithm for 10^5^ iterations with a burn-in of 10^4^. The starting value of ***z*** was set randomly for both algorithms. Each setting of the design was replicated 200 times, and we report Monte Carlo standard errors for all summaries.

### 4.2 Competing methods and metrics

We compare against three alternatives. The first is an ad hoc method that defines a principal balance (Martín-Fernández et al., 2018) from the leading log-ratio principal component, without reference to the mediator or outcome, and then performs product-of-coefficients mediation. The second is regDOC (Wang et al., 2019), a regularized difference-of-coefficients procedure applied to the centered log-ratio transformed taxa. We compare with this method on taxon selection only since it has no comparable definition of an indirect effect. The third is an oracle that fits the model with ***z*** fixed at its true value, providing an upper bound that separates error due to not knowing ***z*** from irreducible noise.

We evaluate performance in estimation of the indirect effect by reporting the bias, root-mean-square error (RMSE), 95% credible-interval coverage, and interval width, together with Type I error at *β* = 0 and power at *β ≠* 0. In addition, we report performance in recovery of the taxa defining the balance, summarized by the true positive rate (TPR) at a false discovery rate (FDR) of 0.1, with the discovery set formed by thresholding the PIPs. Finally, we compare the Gibbs and Metropolis–Hastings samplers of Section 3.3, reporting their per-replicate agreement on the estimated effects and taxon selection, together with mixing (move and acceptance rates), effective samples per CPU-second, and per-iteration computational cost.

### 4.3 Results

Table 1 summarizes estimation of the indirect effect for the proposed method (BalMed), the oracle and the outcome-blind principal balance (PrinBal).

**Table 1:** Indirect-effect estimation under scenarios I & II. The oracle method is given the true support and signs of ***z***—though not the scenario-II weights, whereas the BalMed method needs to infer ***z*** from the data. PrinBal fixes the balance from the leading log-ratio principal component, without reference to the mediator or outcome, and returns a point estimate but no credible interval (its coverage, width, and rejection are therefore omitted, shown as –).

| Scenario | $n$ | $\beta$ | Method | Bias (SE) | RMSE | Cover95 | Width | Reject |
| --- | --- | --- | --- | --- | --- | --- | --- | --- |
| I | 100 | 0 | BalMed | −0.036 (0.003) | 0.057 | 0.730 | 0.134 | 0.270 |
|  |  |  | Oracle | 0.004 (0.002) | 0.034 | 0.960 | 0.132 | 0.040 |
|  |  |  | PrinBal | −0.001 (0.002) | 0.022 | — | — | — |
|  |  | 2 | BalMed | 0.132 (0.012) | 0.214 | 0.585 | 0.450 | 0.965 |
|  |  |  | Oracle | −0.003 (0.008) | 0.115 | 0.925 | 0.406 | 1.000 |
|  |  |  | PrinBal | −0.668 (0.011) | 0.685 | — | — | — |
|  | 200 | 0 | BalMed | −0.001 (0.002) | 0.024 | 0.845 | 0.068 | 0.155 |
|  |  |  | Oracle | 0.001 (0.002) | 0.024 | 0.945 | 0.093 | 0.055 |
|  |  |  | PrinBal | 0.003 (0.001) | 0.021 | — | — | — |
|  |  | 2 | BalMed | 0.016 (0.005) | 0.079 | 0.810 | 0.209 | 1.000 |
|  |  |  | Oracle | −0.001 (0.005) | 0.073 | 0.930 | 0.279 | 1.000 |
|  |  |  | PrinBal | −0.651 (0.010) | 0.667 | — | — | — |
| II | 100 | 0 | BalMed | −0.023 (0.003) | 0.048 | 0.810 | 0.134 | 0.190 |
|  |  |  | Oracle | 0.024 (0.002) | 0.039 | 0.925 | 0.121 | 0.075 |
|  |  |  | PrinBal | −0.001 (0.001) | 0.020 | — | — | — |
|  |  | 2 | BalMed | 0.145 (0.012) | 0.221 | 0.615 | 0.519 | 0.945 |
|  |  |  | Oracle | −0.039 (0.008) | 0.121 | 0.925 | 0.429 | 1.000 |
|  |  |  | PrinBal | −0.674 (0.010) | 0.688 | — | — | — |
|  | 200 | 0 | BalMed | 0.009 (0.002) | 0.027 | 0.795 | 0.070 | 0.205 |
|  |  |  | Oracle | 0.023 (0.002) | 0.032 | 0.830 | 0.086 | 0.170 |
|  |  |  | PrinBal | 0.002 (0.001) | 0.020 | — | — | — |
|  |  | 2 | BalMed | 0.035 (0.006) | 0.091 | 0.780 | 0.224 | 1.000 |
|  |  |  | Oracle | −0.034 (0.005) | 0.082 | 0.920 | 0.295 | 1.000 |
|  |  |  | PrinBal | −0.651 (0.010) | 0.665 | — | — | — |

Under scenario I, the proposed method (BalMed) shows little bias under the null, but its credible intervals are anti-conservative. At *n* = 100, the 95% credible interval achieved 73% coverage, compared with 96% for the oracle, resulting in an inflated Type I error of 27%. Under the alternative, BalMed exhibits a finite-sample bias (0.132, versus *−*0.003 for the oracle) and consequently has larger RMSE and lower coverage, although both methods have high power. Performance improves substantially at *n* = 200: the bias under both the null and alternative were close to zero and the RMSE approaches that of the oracle, although the Type I error of BalMed is still inflated. Scenario II, in which the equal-weight balance is replaced with a variance-matched weighted log contrast, yields broadly similar results, indicating that estimation is relatively robust to misspecification of the balance weights. The oracle also shows some Type I error inflation under scenario II, particularly at *n* = 200, reflecting its use of the correct balance support and signs but misspecified weights. Across both scenarios, the main difference between BalMed and the oracle is uncertainty quantification rather than point estimation, particularly as the sample size increases. While the finite-sample bias at *n* = 100 largely disappears at *n* = 200, the credible intervals remain too narrow, leading to under-coverage and inflated Type I error even when the point estimator is nearly unbiased. The substantially better calibration of the oracle under scenario I suggests that this under-coverage arises primarily from uncertainty associated with estimating the latent balance ***z***. In contrast, PrinBal performs poorly under the alternative across both scenarios and sample sizes, with bias approximately equal in magnitude and opposite in sign to the true indirect effect (*αβ* = 2/3). This occurs because the outcome-blind principal balance captures little of the direction relevant to the mediation pathway.

Figure 3 compares taxon selection for BalMed and regDOC, plotting the TPR against the FDR as the PIP threshold is varied. BalMed outperforms regDOC in both scenarios. At FDR = 0.1, BalMed recovers 57–99% of the active taxa across the grid, against only 9–17% for regDOC; recovery improves with sample size and is slightly lower under the misspecified balance weights (scenario II). These results suggest that BalMed identifies the mediating taxa far more reliably than regDOC, even under misspecification of the balance weights.

**Figure 2.**
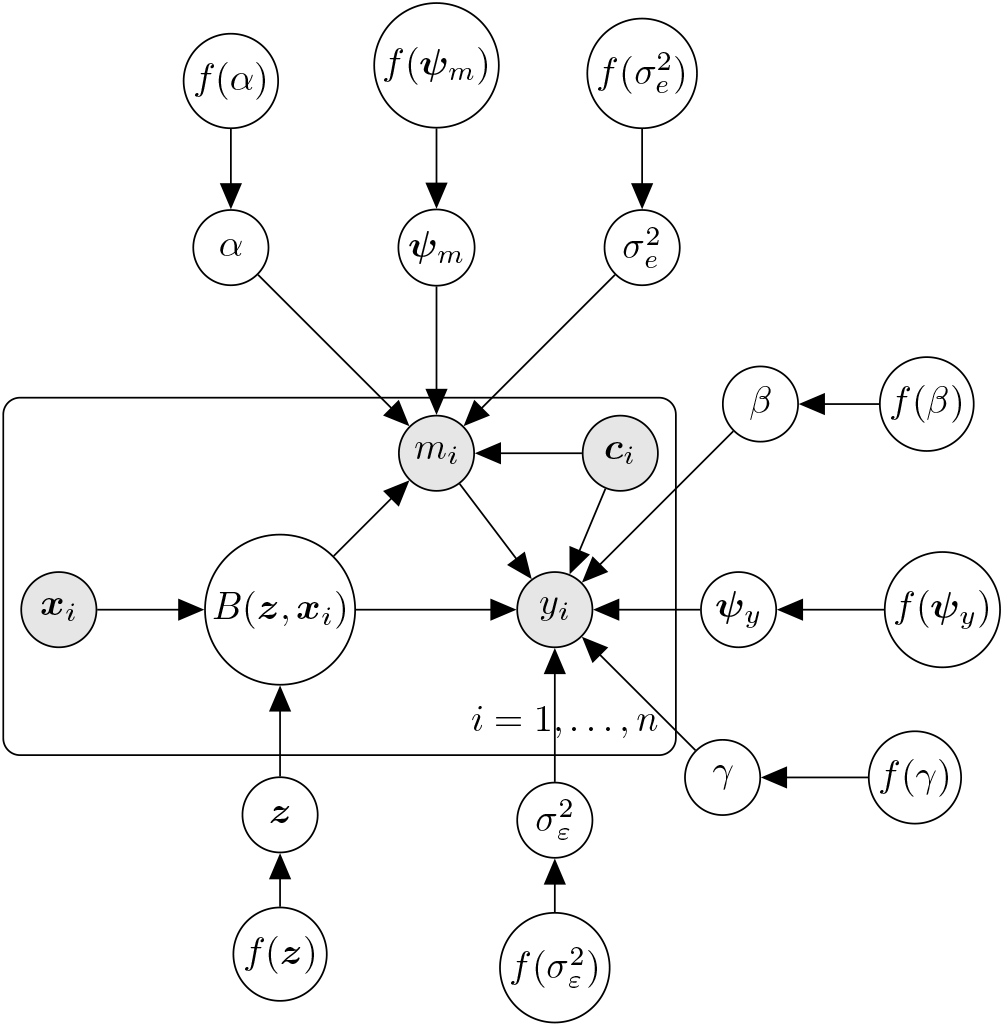
A graphical representation of the proposed mediation framework, where shaded nodes indicate data observed.

**Figure 3.**
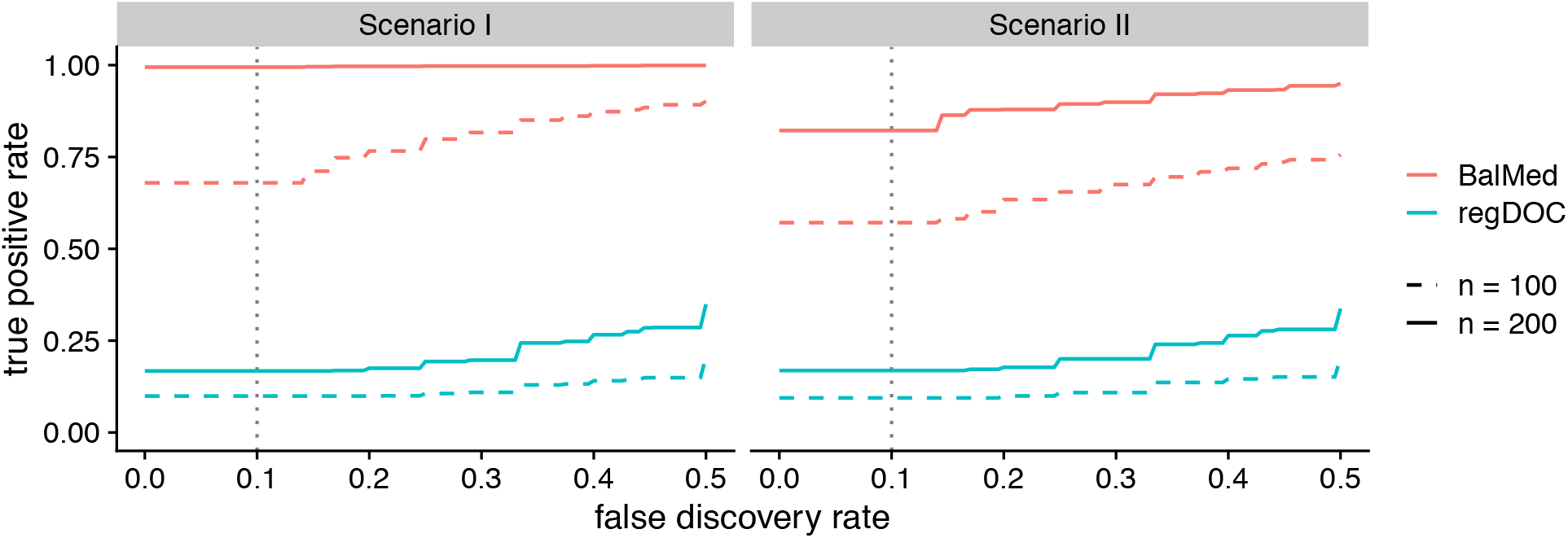
Taxon selection under scenarios I and II: TPR versus FDR for the proposed method and regDOC, by scenario, with method shown by color and sample size by line type (dashed *n* = 100, solid *n* = 200). Each curve is traced by the PIP threshold, averaged over replicates and over *β* (selection is nearly *β*-invariant); the dotted line marks an FDR of 0.1.

Figure 4A–C examines sensitivity to three design factors varied one at a time from scenario I: the prior on ***z*** (panel A, *η*_1_ = *η*_2_ *∈ {*1/3, 1/5, 1/20*}*), the number of taxa (panel B, *d ∈ {*30, 50, 100*}*), and the number of active taxa in each group (panel C, *a ∈ {*1, 3, 5, 10*}*). Solid and dashed lines correspond to the null and non-null mediator settings, respectively. A sparser prior modestly improves taxon selection, with TPR at FDR = 0.1 increasing from 0.60 to 0.76 as *η* decreases from 1/3 to 1/20), along with a reduction in indirect-effect RMSE. Performance declines with increasing *d*, particularly at *d* = 100 where the number of taxa equals the sample size. Taxon recovery also deteriorates as the true balance becomes less sparse, with TPR decreasing from 0.89 at *a* = 1 to 0.37 at *a* = 10, although the indirect-effect estimation is comparatively stable.

**Figure 4.**
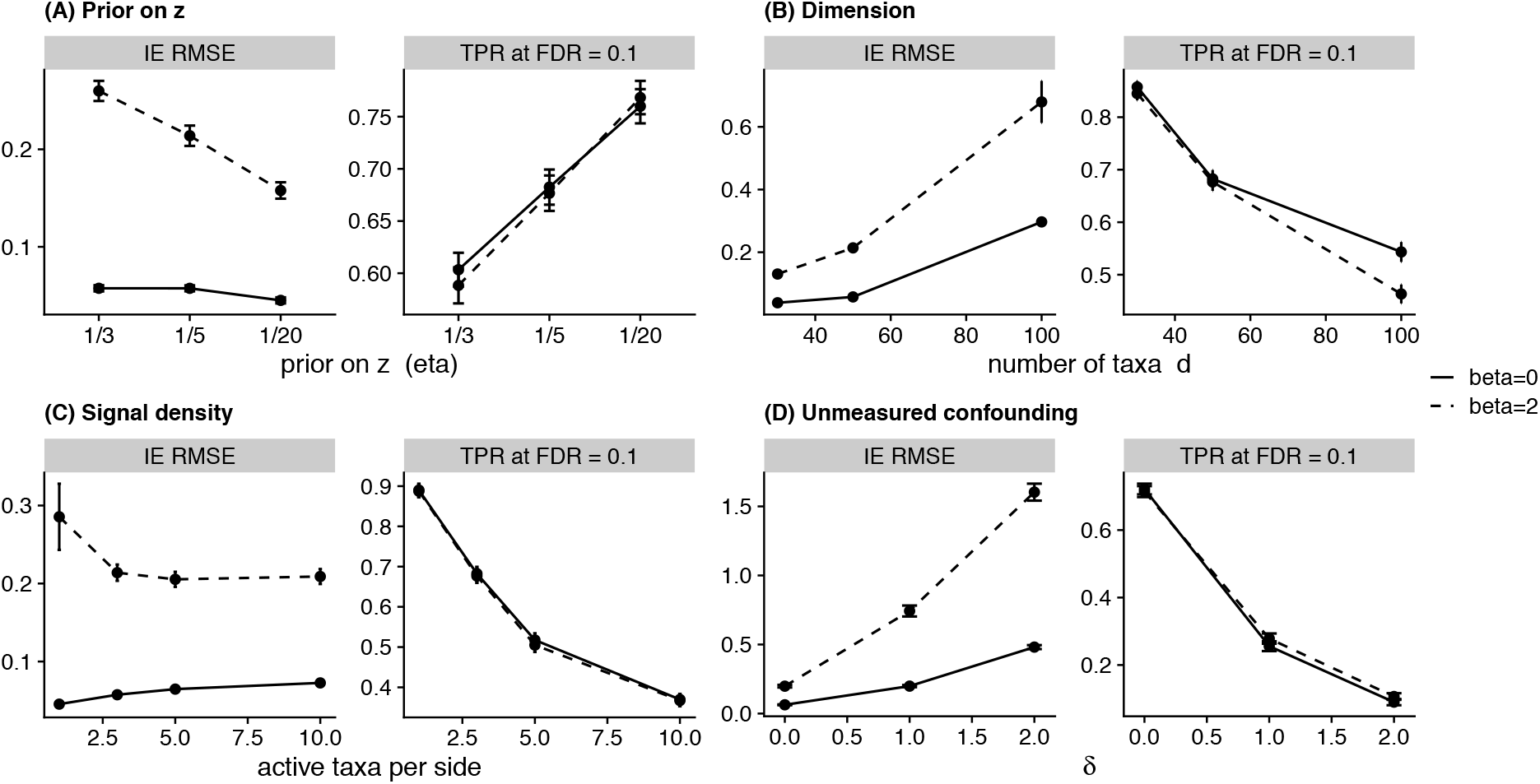
One-factor sensitivity and robustness analyses for the proposed method. Panels vary the prior on ***z*** (A), the number of taxa *d* (B), the number of active taxa per side *a* (C), and the confounder loadings *δ*_*m*_ = *δ*_*y*_ = *δ* (D). Each panel reports indirect-effect RMSE (left) and the TPR at FDR = 0.1 (right). Solid lines are the null mediator (*β* = 0) and dashed lines the non-null (*β* = 2); bars are Monte Carlo standard errors.

Figure 4D examines sensitivity to unmeasured mediator–outcome confounding under scenario III, in which an unobserved variable *u*_*i*_ affects both the mediator and the outcome with *δ*_*m*_ = *δ*_*y*_ = *δ ∈ {*0, 1, 2*}* but is omitted at estimation. As expected from the identification assumptions, indirect-effect estimation is accurate at *δ* = 0 but deteriorates as the strength of confounding increases. Taxon selection similarly worsens with increasing *δ*, indicating that the unmeasured mediator–outcome confounding affects both estimation of indirect effects and recovery of the latent balance. These results motivate caution in interpreting the mediation effects in the data application.

Finally, Figure 5 compares the Gibbs and MH samplers under a common simulation setting. Estimates of indirect and direct effects from the two samplers largely agree, whereas agreement on the taxon-selection TPR at FDR = 0.1 is weaker (panel A). Across replicates the Gibbs–MH differences are centered near zero but are more variable for taxon selection than for the effect estimates, which is consistent with the greater sensitivity of threshold-based selection summaries to Monte Carlo variation. The Gibbs sampler is substantially more efficient despite its higher per-iteration cost (panel B). A Gibbs iteration costs about 1.5 times as much CPU as an MH iteration, but explores the balance space more effectively, with a move-rate approximately 23 times higher than the MH acceptance rate, which is below 4%. Consequently, Gibbs yields approximately 5.7 times as many effective samples per CPU-second. A Gibbs sweep costs approximately 0.23 seconds on a single core of an Intel Xeon Gold 6254 processor (3.10 GHz). These results support Gibbs as the default sampler, with MH providing an alternative when the computational cost of Gibbs becomes prohibitive at large *d*.

**Figure 5.**
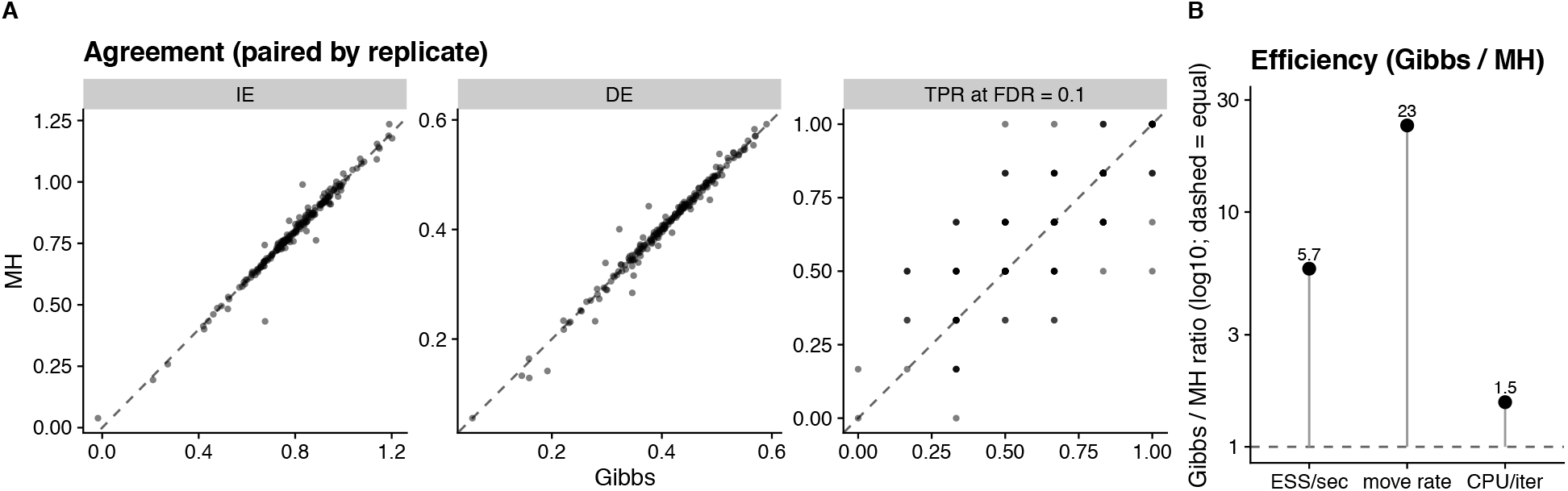
Gibbs versus MH on a common simulation setting (*n* = 100, scenario I, *β* = 2). (A) Agreement: each point is one replicate, plotting the MH value against the Gibbs value for the indirect effect (IE), direct effect (DE), and the taxon-selection TPR at FDR = 0.1; the dashed line is *y* = *x*. (B) Efficiency, as Gibbs-to-MH ratios (log scale) of effective samples per CPU-second, move rate, and CPU per iteration; the dashed line is *y* = 1. These ratios are per-second or per-iteration, so they are unaffected by the different run lengths.

## 5 Application

We illustrate the proposed method using data from the MEC-APS study (Hullar et al., 2021). Between 2013 and 2016, MEC-APS characterized the fecal microbiome by 16S rRNA gene sequencing in healthy older adults from five self-reported racial and ethnic groups, and collected magnetic-resonance measures of hepatic fat together with demographic and clinical variables. Using these data, we examine whether the association between the gut microbiome and percent liver fat is mediated by lipopolysaccharide-binding protein (LBP). LBP is an acute-phase protein that binds lipopolysaccharide (LPS) released by Gram-negative gut bacteria and is commonly used as a marker of metabolic endotoxemia, a potential mechanism linking the gut microbiome to hepatic fat accumulation and insulin resistance (Cani et al., 2007; Moreno-Navarrete et al., 2012). We take the microbiome, represented by the latent balance *B*(***z, x***_*i*_), as the exposure; LBP as the mediator *m*_*i*_; and percent liver fat as the outcome *y*_*i*_.

The analytic sample comprises *n* = 1513 participants with complete mediator, outcome, and covariate information, including metformin status. We retain genera with relative abundance at least 0.01% in 10% or more of the samples, resulting in 135 genera. Zeros, which comprise about 17% of the filtered abundance matrix, are replaced by a pseudocount of 0.5. All models adjust for age, sex, self-reported race and ethnicity, place of birth, education, total body-fat percentage, the Alternative Healthy Eating Index–2010 diet-quality score, smoking pack-years, sedentary time, and season of stool collection. We adjust for measured total body-fat percentage rather than body mass index because the latter is an imperfect surrogate for adiposity whose accuracy differs across racial and ethnic groups (Hullar et al., 2021). Furthermore, because diabetes is associated with higher liver fat and systemic inflammation (Forslund et al., 2015), the primary model additionally adjusts for diabetes status. In this cohort, diabetes and metformin use are nearly coextensive (162 of the 192 metformin users have diabetes), and because both may be downstream of the microbiome rather than pre-exposure confounders (Aron-Wisnewsky et al., 2020), we also fit models that replace the diabetes indicator with an indicator of metformin use, or omit both. Given the cross-sectional design, causal interpretation of the estimated direct and indirect effects relies on the identification assumptions in Section 2, whose plausibility we revisit in Section 6.

Figure 6 reports the effect decomposition (left) and path coefficients (right) for the primary model, which uses the sparse prior *η* = 1/20 and adjusts for diabetes status, together with seven sensitivity fits. In the primary model the direct effect is positive (*γ* = 0.12, 95% credible interval 0.10 to 0.13), as is the exposure–mediator coefficient (*α* = 0.031, 95% CrI 0.021 to 0.041). The mediator–outcome coefficient is also positive (*β* = 0.075, 95% CrI 0.012 to 0.139), giving a small indirect effect (*αβ* = 0.0023, 95% CrI 0.0004 to 0.0045) that amounts to 2.0% (95% CrI 0.3% to 3.8%) of the total effect. The direct effect and *α* are credibly positive across all eight models, but the indirect effect is less certain. Across the seven models that adjust for adiposity its posterior mean ranges from 0.0017 to 0.0030 and the posterior probability that it is positive is at least 0.96, but the 95% credible interval includes zero under the less sparse priors (*η* = 1/3, 1/5) and in the two exclusion fits, where metformin users or participants with diabetes are dropped. Omitting the adiposity covariate roughly doubles both *β* (0.16, 95% CrI 0.09 to 0.22) and the indirect effect (0.0053, 95% CrI 0.0030 to 0.0079). This contrast suggests that much of the LBP–liver-fat association is shared with adiposity, and that the component that remains after conditioning on adiposity is small. Mediation operating through adiposity itself, for example if endotoxemia raises liver fat by promoting overall adiposity, would be absorbed by the adjustment and cannot be separated from the direct effect with these data. All chains converge 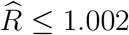 for every reported quantity), and taxon-inclusion probabilities agree closely across chains (minimum pairwise cross-chain correlation 0.998). Convergence diagnostics for the primary model are presented in Appendix B.

**Figure 6.**
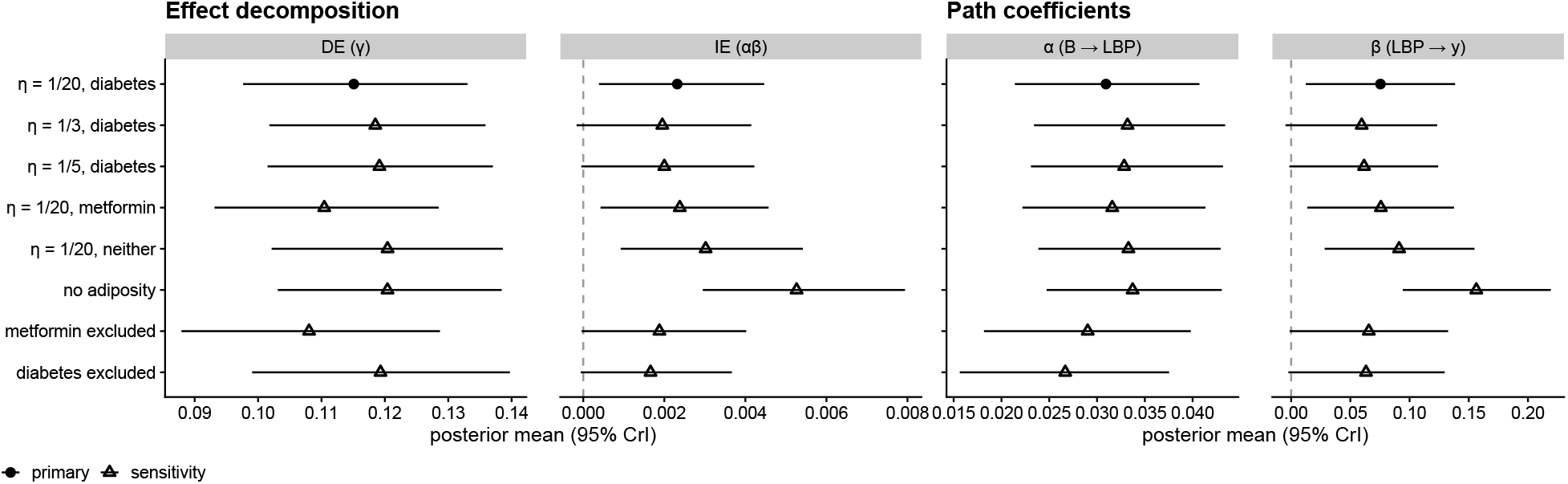
Effect decomposition (left) and path coefficients (right) for the MEC-APS analysis, estimated by the Gibbs sampler. The primary model (filled circle) uses the sparse prior *η* = 1/20 and adjusts for diabetes status (*n* = 1513, metformin users retained). Open triangles are sensitivity fits, each altering the primary model in exactly one respect: the prior (*η ∈* {1/3, 1/5}), the additional covariate (metformin use, or neither, in place of diabetes status), the adiposity adjustment (omitted), or the sample (excluding metformin users, *n* = 1321; excluding participants with diabetes, *n* = 1266, which then adjusts for neither diabetes nor metformin). Points are posterior means and horizontal bars are 95% credible intervals. Dashed lines mark zero in the *αβ* and *β* panels; the *γ* and *α* axes are restricted to the range of the estimates, which lie entirely above zero.

Lastly, we threshold the component-wise PIPs at 0.5 to select taxa that characterize the latent balance. Table 2 reports the 21 genera selected in the primary model; 10 of them are selected in all eight model fits shown in Figure 6, indicating a stable core set of taxa. Under the sign convention *γ >* 0 and *α >* 0, higher abundance of the denominator taxa relative to the numerator is associated with lower liver fat through the direct path and with lower circulating LBP. The denominator consists largely of obligate anaerobes of the Oscillospiraceae, Ruminococcaceae, and Lachnospiraceae, including *Oscillospira, Intestinimonas, Anaerotruncus*, and the Lachnospiraceae NK4A136 group, several of which are short-chain fatty-acid producers (Koh et al., 2016), in line with the depletion of butyrate-producing bacteria reported in fatty liver disease (Aron-Wisnewsky et al., 2020). It also contains *Haemophilus*, a Gram-negative genus whose lipooligosaccharide is a ligand of LBP; because the balance is a joint summary of co-varying taxa, the side of any single taxon should be interpreted as its contribution to circulating LBP relative to the others in the balance. The numerator comprises the Gram-negative genera *Bacteroides, Veillonella*, and the bile-tolerant sulfite reducer *Bilophila*, several Lachnospiraceae (*Howardella, Lachnoclostridium, Marvinbryantia*, CAG-56, and an uncultured Lachnospiraceae clade), *Colidextribacter, Negativibacillus*, and *Candidatus* Stoquefichus. Because the direct effect is an order of magnitude larger than the indirect effect, the selected balance is identified primarily by the direct microbiome–liver-fat association.

**Table 2:** The 21 genera selected (PIP *>* 0.5) in the primary model (*η* = 1/20, diabetes-adjusted). Signed inclusion is Pr(*z*_*j*_=+1) − Pr(*z*_*j*_=−1); a positive value places the taxon in the numerator of the balance and a negative value in the denominator, and its magnitude reflects the selection frequency. The last column counts the models in Figure 6 (of eight) in which the taxon is selected. Placeholder labels carry the parent family from the study taxonomy; “uncultured” denotes sequence variants assigned to the family but to no named genus.

| Taxon | Signed inclusion | Models (of 8) |
| --- | --- | --- |
| Lachnospiraceae (family; uncultured) | +1.00 | 8 |
| Marvinbryantia | +0.99 | 8 |
| Lachnoclostridium | +0.96 | 8 |
| Veillonella | +0.91 | 6 |
| Bacteroides | +0.89 | 8 |
| Lachnospiraceae CAG-56 | +0.76 | 8 |
| Howardella | +0.74 | 6 |
| Colidextribacter | +0.73 | 6 |
| Negativibacillus | +0.67 | 6 |
| *Candidatus* Stoquefichus | +0.60 | 7 |
| Bilophila | +0.60 | 7 |
| Clostridia UCG-014 | −1.00 | 8 |
| Oscillospira | −1.00 | 8 |
| Erysipelatoclostridium | −0.97 | 8 |
| Intestinimonas | −0.95 | 8 |
| Haemophilus | −0.90 | 6 |
| Lachnospiraceae NK4A136 group | −0.89 | 8 |
| Christensenellaceae (family; uncultured) | −0.82 | 7 |
| Anaerotruncus | −0.66 | 7 |
| Holdemania | −0.65 | 5 |
| Cloacibacillus | −0.51 | 5 |

## 6 Discussion

We propose a Bayesian framework for mediation analysis with a compositional exposure. The high-dimensional microbial composition is summarized by a latent balance, a log-ratio between two unknown groups of taxa, that acts as a scalar exposure driving both the mediator and the outcome, so that the community-level direct and indirect effects reduce to simple functions of a single log-ratio while the taxa defining the balance are selected as part of the model. Estimation and taxon selection are performed jointly through a collapsed Gibbs sampler, with a Metropolis–Hastings alternative for very high dimensions. In simulations, the method estimates the indirect effect and recovers the balance-defining taxa accurately, remains robust to misspecification of the balance shape, and, as the identification theory predicts, is sensitive to unmeasured mediator–outcome confounding. When applied to the MEC-APS cohort, it summarizes the gut microbiome by an interpretable balance that has a robust direct association with hepatic fat, and finds a small indirect effect through LBP, whose credible interval excludes zero in the primary model but not under every sensitivity setting.

The MEC-APS estimates are observational and should be interpreted under the identification assumptions of Section 2. The stable unit treatment value assumption and positivity are plausible for a continuous latent exposure, but the two consequential assumptions—no unmeasured confounding and correct temporal ordering—are not guaranteed by the cross-sectional design. Unmeasured factors such as hepatic inflammation and host genetics could confound the LBP–liver-fat relationship, and the assumed ordering microbiome *→* LBP *→* liver fat cannot be verified. Our simulations show that the estimated indirect effect is sensitive to precisely this mediator–outcome confounding, which can bias the estimate in either direction. The small indirect effect should therefore be interpreted as weak evidence for an adiposity-independent LBP pathway under the stated assumptions rather than as an established mechanism; that the estimate roughly doubles when adiposity is not adjusted for shows how much it depends on the covariate set, and confirmatory analysis would require longitudinal or interventional data.

This work can be extended in several directions. The present paper focuses on a single mediator, which matches the data application. An extension to multiple, possibly correlated, mediators can be achieved by generalizing the mediator model to a free residual covariance and adapting the coordinated selection framework of Song et al. (2021) to select active mediators jointly with the balance. In addition, the current framework assumes the direct and indirect effects of the microbiome on the outcome are through a single latent log contrast. While this simplifies interpretation, it does impose the assumption that the effect of the microbiome is explained via a sparse set of taxa which collectively define the balance. When the effects of individual taxa are more diffused, this assumption may be violated. In those settings, auxiliary information, such as the phylogenetic tree, may provide alternative structural constraints such that the mediation effects are still identifiable.

## Data Availability Statement

The data from the Multiethnic Cohort–Adiposity Phenotype Study that support the findings of this study are available upon reasonable request. Code implementing the proposed method and reproducing the simulation studies is available at https://github.com/drjingma/BalExMed-paper. An R package implementing the proposed method is available at https://github.com/drjingma/BalExMed.

## Acknowledgements

This work is supported by NIH grant GM145772.

## A Derivations of the MCMC algorithm

Let 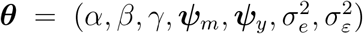 denote the list of parameters in the model. The joint likelihood *f* (***y, m*, X, C, *θ*** | ***z***) can be written as the product of two terms *L*_1_*L*_2_, where

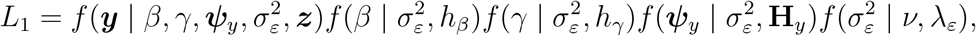

and

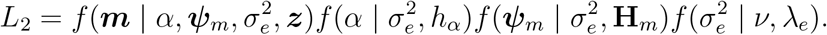

We first derive the conditional density of ***y*** given ***m*, X, C** and ***z***. Let ***ζ*** = (*β, γ*, ***ψ***_*y*_) denote all the coefficients in the outcome regression model and **H**_1_ = *diag*(*h*_*β*_, *h*_*γ*_, **H**_*y*_). Recall **D**_*y*_ = [***m*, B**_*z*_, **C**] is the design matrix in the outcome regression model. The outcome regression model can be rewritten as

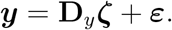

The joint likelihood *L*_1_ is

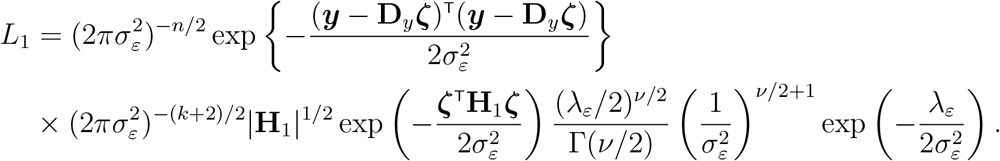

Let 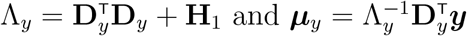. We can rewrite the joint likelihood as

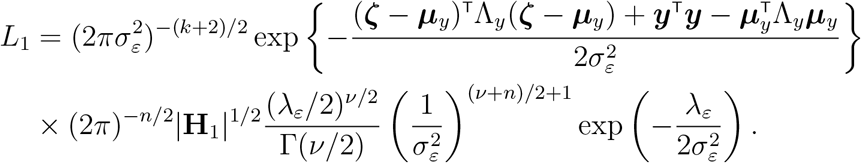

It is easy to see that the posterior of ***ζ*** given 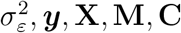 and ***z*** is multivariate normal with mean ***µ***_*y*_ and variance 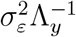. The posterior density of 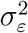 given ***y*, X, M, C** and ***z*** is

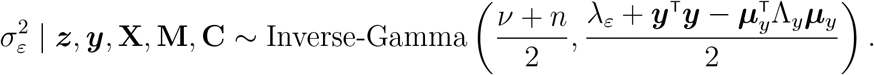

We can obtain the conditional density ***y*** given ***z*** by integrating out ***ζ*** and 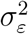

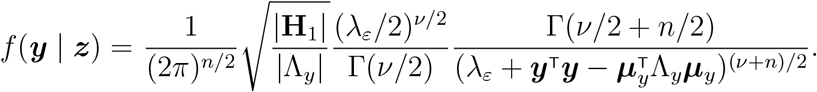

To derive *L*_2_, noting that 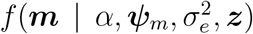 is the density of a multivariate normal distribution with mean *α***B**_*z*_ + **C*ψ***_*m*_ and variance 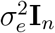, we have

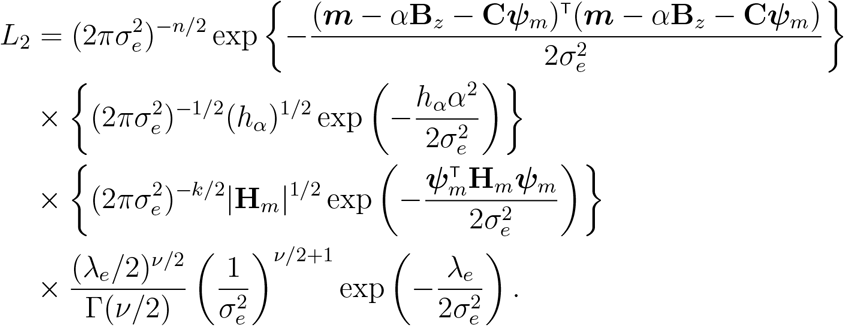

Let 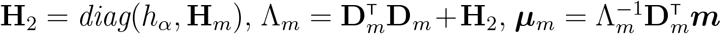 and ***ξ*** = (*α*, ***ψ***_*m*_). Rearranging terms, we can rewrite *L*_2_ as

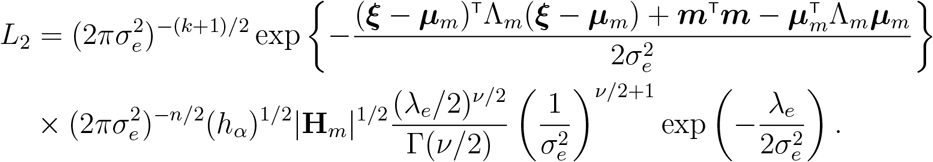

It is easy to see that the posterior of ***ξ*** given 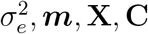 and ***z*** is multivariate normal with mean ***µ***_*m*_ and variance 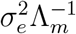. The posterior distribution of the variance parameter 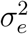 is

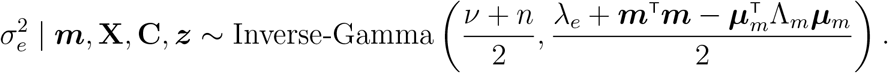

The normal-inverse-gamma posterior allows us to integrate out the parameters and obtain the conditional density

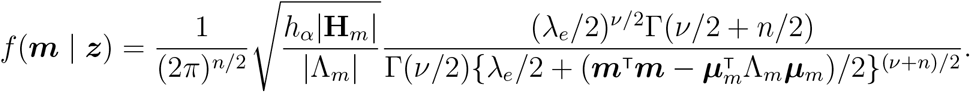

Putting all pieces together, we have the conditional density of the observed data given the selection index ***z***

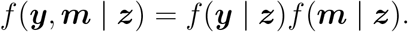

Additionally, the conditional density of the coefficients ***ζ*** and ***ξ*** given the selection index ***z*** can be derived after integrating out the variance components. Specifically, ***ζ*** | ***z, y, m*, X, C** is a multivariate *t*-distribution with degrees of freedom *ν* + *n*, location parameter ***µ***_*y*_, and shape parameter

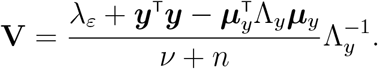

Similarly, ***ξ*** | ***z, m*, X, C** has a *t*-distribution with degrees of freedom *ν* + *n*, location parameter ***µ***_*m*_, and shape parameter

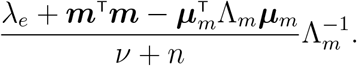

## B Convergence diagnostics for the MEC-APS analysis

Every MEC-APS fit uses four chains of 5000 iterations, of which the first 1000 are discarded as burn-in. Chain 1 starts from the leading principal component of the centered log-ratio transformed genera abundances and the other three start from random configurations, so the chains begin far apart in the space of balance configurations. We report the diagnostics for the primary model. The seven sensitivity fits behave similarly: across all eight fits the largest split-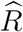 is 1.002, the smallest effective sample size is 3965, and the smallest pairwise correlation between the taxon-inclusion probabilities of two chains is 0.998.

Figure S1 shows the traces and running means of the direct effect *γ* and the indirect effect *αβ*. The four chains explore the same region from the first retained draw, and their running means coincide within a few hundred draws. All split-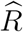 values are at most 1.001. The effective sample size is smallest for *α*, at about 6000 of the 16,000 retained draws, and largest for the indirect effect, at about 15,500.

**Figure S1.**
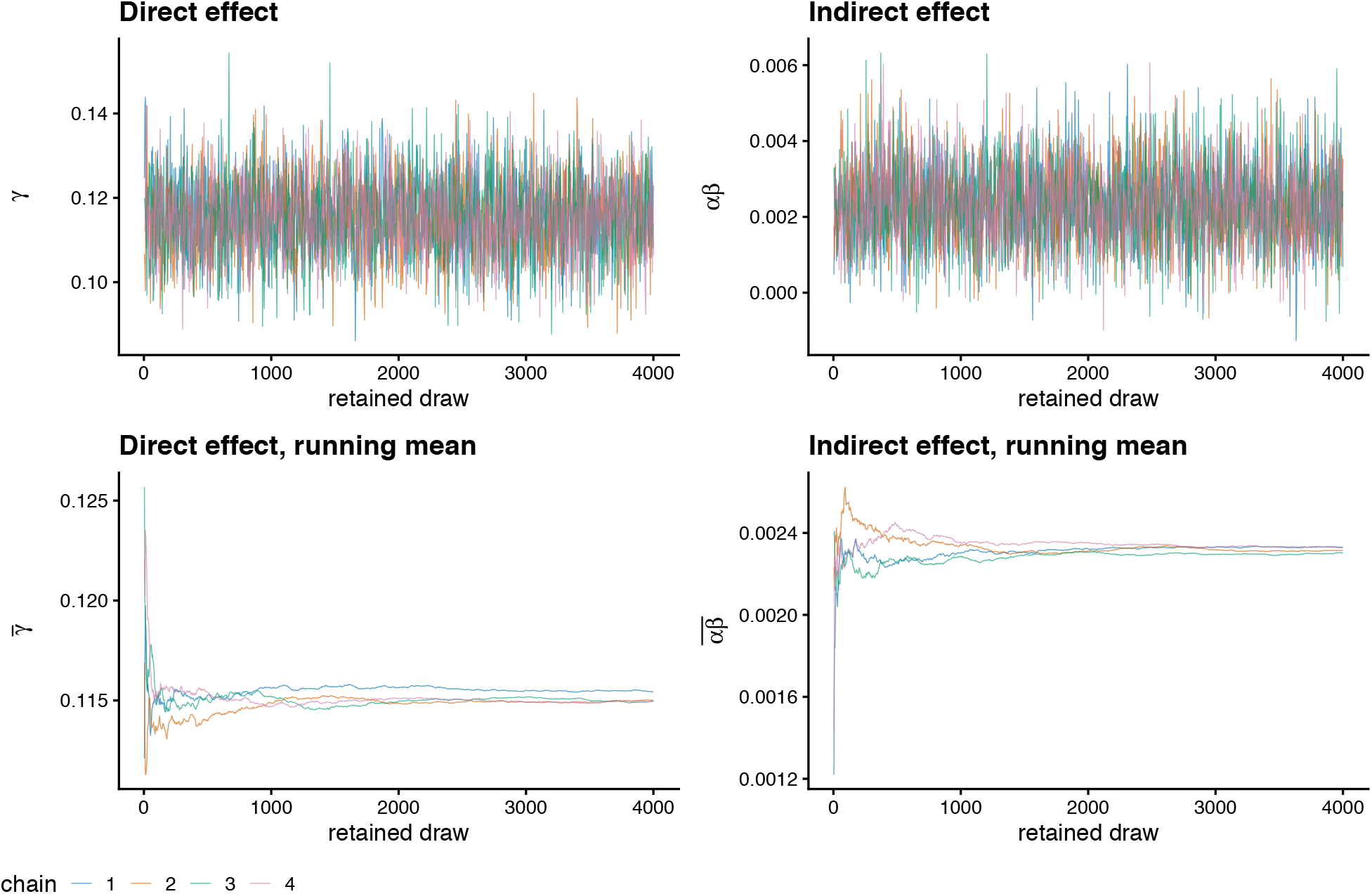
Convergence of the primary MEC-APS fit (*η* = 1/20, diabetes-adjusted). Traces of the direct effect *γ* and the indirect effect *αβ* (top) and their running means (bottom), with one colour per chain. Every fourth retained draw is shown; the running means use every draw.

Figure S2 examines mixing over the balance configuration ***z***, which the scalar summaries above do not capture. Panel A compares the posterior inclusion probability of each genus, computed separately within each chain, against the values from chain 1. The points lie along the diagonal: the smallest pairwise correlation between chains is 0.999 and the largest across-chain standard deviation of an inclusion probability is 0.033. Panel B shows how many of the 135 coordinates of ***z*** change in each Gibbs sweep. After burn-in the sampler changes a median of 9 coordinates per sweep and changes at least one coordinate in at least 99.9% of sweeps, so the chains keep moving between configurations rather than settling on a single one. The relabelling step that enforces *γ >* 0 is triggered in 0.02% of iterations, consistent with a direct effect that is well separated from zero.

**Figure S2.**
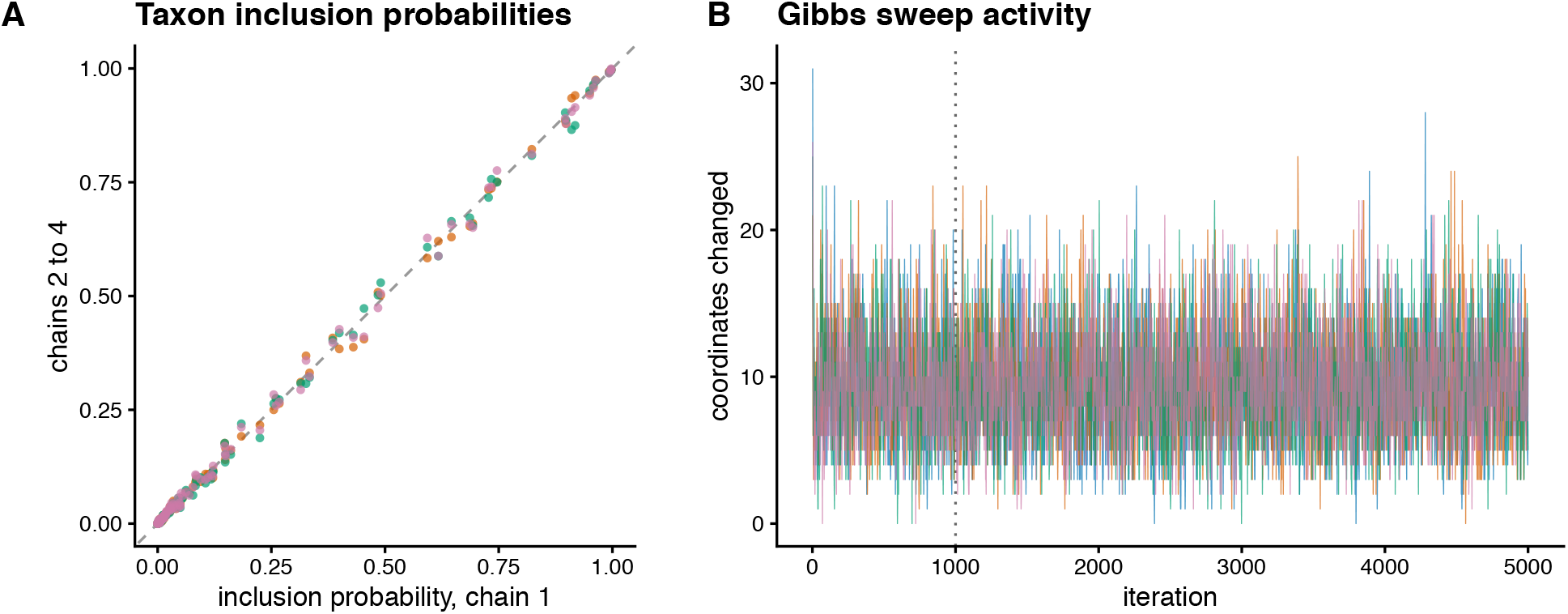
Mixing across chains in the primary MEC-APS fit. (A) Posterior inclusion probability of each genus in chains 2 to 4 against the corresponding value in chain 1; the dashed line is *y* = *x*. (B) Number of coordinates of the balance configuration ***z*** changed in each Gibbs sweep, one line per chain and every second iteration shown; the dotted vertical line marks the end of burn-in.

